# Distinct function of Mediator subunits in fungal development, stress response and secondary metabolism in fungal pathogen *Fusarium verticillioides*

**DOI:** 10.1101/2021.02.11.430821

**Authors:** Zehua Zhou, Huijuan Yan, Man S. Kim, Won Bo Shim

## Abstract

Mediator is a conserved, nucleus-localized, multi-subunit complex highly conserved across eukaryotes. It interacts with RNA polymerase II transcription machinery as well as various transcription factors to regulate gene expression. However, systematic characterization of Mediator complex has not be thoroughly performed in filamentous fungi. In current study, our aim was to investigate key biological functions of Mediator subunits in a mycotoxigentic plant pathogen *Fusarium verticillioides*. While we recognized some level of divergence in subunits constituent, overall structure remained conserved between *Saccharomyces cerevisiae* and *F. verticillioides*. We were able to generate 11 Mediator subunit deletion mutants and characterize serveal important phenotypes associated with fungal development, environmental stress, nutrient utilization, virulence as well as secondary metabolism. While each Mediator subunit deletion mutant showed deficiencies in at least three of the phenotypes tested, it is important to note that different modules or subunits showed unique regulatory role in different cellular processes. Significantly, the deletion of *FvMed1* led to increased FB_1_ production, and we were able to confirm that FvMed1 is transported from the nucleus to the cytoplasm under fumonisin-producing conditions. Taken together, our study characterized the different biological functions of Mediator subunits in plant pathogen *F. verticillioides* and the possibility of select subunits unique cytoplasmic function independent of the core Mediator.

**IMPORTANCE:** Mediator is a highly conserved component of RNA polymerase II pre-initiation complex (PIC) in eukaryotes. Studies in yeast have demonstrated the important roles of Mediator subunits, but we lack deeper understanding in filamentous fungi. Here, we selected *Fusarium verticillioides*, a major fungal pathogen of maize worldwide, to study transcriptional profile followed by mutational analyses of all Mediator genes. Each Mediator subunit deletion mutant showed some shared deficiencies but it is also important that different modules or subunits showed unique regulatory role in different cellular processes. Significantly, *FvMed1* deletion mutant showed elevated levels of FB_1_ production, and we were able to confirm that FvMed1 translocates from the nucleus to cytoplasmic organelles under fumonisin-producing conditions. This is a first study demonstrating that a Mediator subunit regulate cellular functions while not directly associating with PIC in filamentous fungi.

## INTRODUCTION

Cellular growth, differentiation and development is a complex and highly regulated process that relies on differentially and methodically coordinated expression of numerous genes in fungi. The coordinated expression, including protein-coding genes and various non-coding RNAs, is accomplished by the RNA polymerase II (Pol II) complex. Notably, Pol II holoenzyme does not carry out transcription alone but performs function as a macromolecular structure known as the pre-initiation complex (PIC), which consist of Pol II, Mediator, TFIIA, TFIIB, TFIID, TFIIE, TFIIF and TFIIH (1, 2). Among these components, Mediator serves as the central scaffold of PIC and is known as the main regulator of Pol II activity. However, owing to its structural complexity and expansive size, our understanding of the biochemical and genetic functions of Mediator in filamentous fungi is rudimentary.

Mediator complex was first discovered in *Saccharomyces cerevisiae* (3), and their organization and structure seem to be evolutionarily conserved among eukaryotes. Mediator complex contains 30 (human) or 25 (yeast) subunits which can be classified into four functional modules: Head, Middle, Tail and Kinase. Significantly, the subunit 14 (Med14) serves as a scaffold and holds the Head, Middle and Tail modules together as a complex, which is defined as the ‘core Mediator’ (4–6). Published studies indicated that the ‘core Mediator’ serves as a bridge between DNA-binding transcription factors (TFs), Pol II as well as other components in the transcription machinery, which collectively plays a crucial role in transcriptional activation (7–10). The Kinase module, which consists of four subunits Med12, Med13, Cdk8 (cyclin-dependent kinase 8) and Ssn (cyclin C), could associate reversibly with the ‘core Mediator’, thereby regulating Pol II activity and transcription (11–13).

While the functions of Mediator appear highly conserved among eukaryotes, analyses of Mediator mutants in various cells and tissues have shown that each subunit of the Mediator complex regulates distinct subsets of genes during various developmental processes. For example, Ssn3 and Ssn8 are required for glucose metabolism and biofilm formation in *C. albicans* (14). In *S. cerevisiae*, Med5 is involved in expression of genes associated with nuclear and mitochondrial oxidative phosphorylation (OXPHOS) (15). It is interesting to note that while Med7 is essential for yeast viability, its ortholog in *C. albicans* is not essential and is involved crucial developmental control (16). Med31 is a maternal-associated gene in *Drosophila* (17), whereas no defects have been detected during early embryonic growth in mouse mutants (18). These reports support gene- and species-specific functions within Mediator even though it has many conserved roles across eukaryotes. Although the biological and physiological functions have been extensively studied in a select number of model organisms, little is known about Mediator functions filamentous fungi.

*Fusarium verticillioides* is the fungal pathogen responsible for devastating stalk rot, ear rot and seedling blight diseases in maize, which can lead to severe yield losses worldwide (19, 20). Importantly, *F. verticillioides* produces a variety of mycotoxins during maize infestation in the field or in storage. Among these mycotoxins, fumonisins B1 (FB1) poses a particular concern, with links to birth defects and cancer in humans as well as acute and chronic toxicosis in animals (21–24). Therefore, the concentration of fumonisins in maize and food product is regulated by governmental agencies worldwide (25). One of the key challenges in developing effective mitigation strategies for fumonisins in food and feeds is our limited understanding of how fumonisin biosynthesis is regulated in *F. verticillioides*. Published studies showed that Kinase module subunits, namely cyclin C and cyclin C-dependent kinase, play a crucial role in mycotoxin biosynthesis in *Fusarium* species (26–29), but the role of other Mediator subunits is uncharacterized to date.

In order to investigate the functions of Mediator subunits, we implemented a systematic study of all Mediator subunit genes in *F. verticillioides* by transcriptional analyses and gene knockout strategy. In our study, 23 Mediator subunit genes were found in *F. verticillioides*, and the functional roles of 11 Mediator subunits were investigated in this study. Our results provide further evidence that different modules and subunits play a unique role in transcriptionl regulation of specific subsets of genes in *F. verticillioides*. Significantly, the translocation of a Mediator subunit from the nucleus to the cytoplasm was observed in fumonisin-producing conditions and non-producing conditions, suggesting that Mediator subunits may perform yet-to-be determined functions in the cytoplasm independent of Pol II.

## RESULTS

### Mediator subunit composition in *S. cerevisiae* and F. *verticillioides*

In all eukaryotes, including *S. cerevisiae*, Mediator subunits have been categorized into four functional modules: Head, Middle, Tail and Kinase (4, 15, 30, 31). We used *S. cerevisiae* subunits as queries and screened *F. verticillioides* predicted proteins for putative orthologs. Table 1 describes how Mediator subunits in two fungi can be comparatively organized. Although the Mediator subunit organization is conserved between *F. verticillioides* and *S. cerevisiae*, we did learn that *S. cerevisiae* subunits Med20, Med10, Med19 and Med2 were not identifiable in *F. verticillioides*. Also, we found that *F. verticillioides* harbors 15 copies of *FvMED15* gene while the gene encoding *MED15* resides as a single copy locus in *S. cerevisiae*. These genes are named as the telomere-associated genes *(TLOs)* and may suggest the evolutional divergence of Med15 between *F. verticillioides* and *S. cerevisiae*.

**Table 1.**
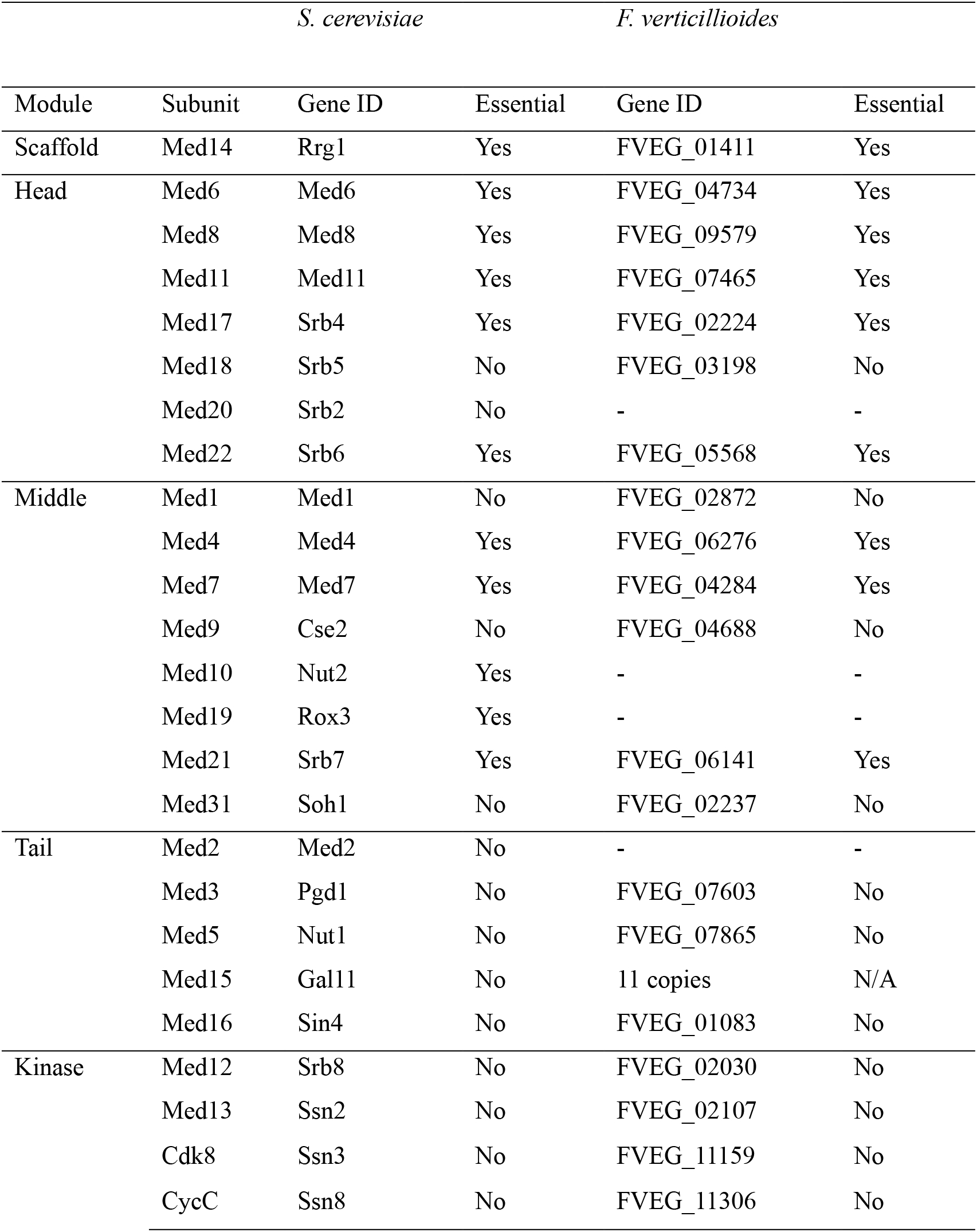
Comparisons of Mediator modular organization and subunit composition between *Fusarium verticillioides* and *Saccharomyces cerevisiae*.

### Transcription profile of Mediator genes

To determine whether certain Mediator subunits are transcriptionally associated with FB1 biosynthesis regulation, we analyzed their expression levels in RNA-Seq libraries generated from *F. verticillioides* cultured in maize inbred line B73 kernels (moderate FB1 production) versus hybrid 33K44 kernels (high FB1 production). We followed the relative transcription levels of Mediator subunits in 2, 4, 6, 8 days post inoculation (dpi) samples (32). Overall, the expression of each Mediator gene increased through the time course and highest level at 8 dpi, especially noticeable for *FvMed14 (FVEG_01411), FvMed12 (FVEG_02030), FvMed31 (FVEG_02237), FvMed7 (FVEG_04284), FvMed21 (FVEG_06141)* and *FvMed3 (FVEG_07603)* (Fig. S1). However, it is important to note that the expression of all Mediator genes showed no statistical difference between two RNA-Seq libraries as shown in Fig. 1 where we highlighted key Head *(FVEG_03198)*, Middle *(FVEG_02872, FVEG_02237)*, Tail *(FVEG_07603)* and Kinase *(FVEG_11159, FVEG_11306)* subunits. Notably, transcription profile of cyclin C *FvFcc1 (FVEG_11306)* and Cyclin-dependent kinase *FvFck1 (FVEG_11159)* were in agreement with earlier published studies (26, 27). These outcomes led us to predict Mediator subunits do not regulate FB1 biosynthesis transcriptionally but rather by forming complex with core Mediator or PIC in *F. verticillioides*.

**Fig. 1.**
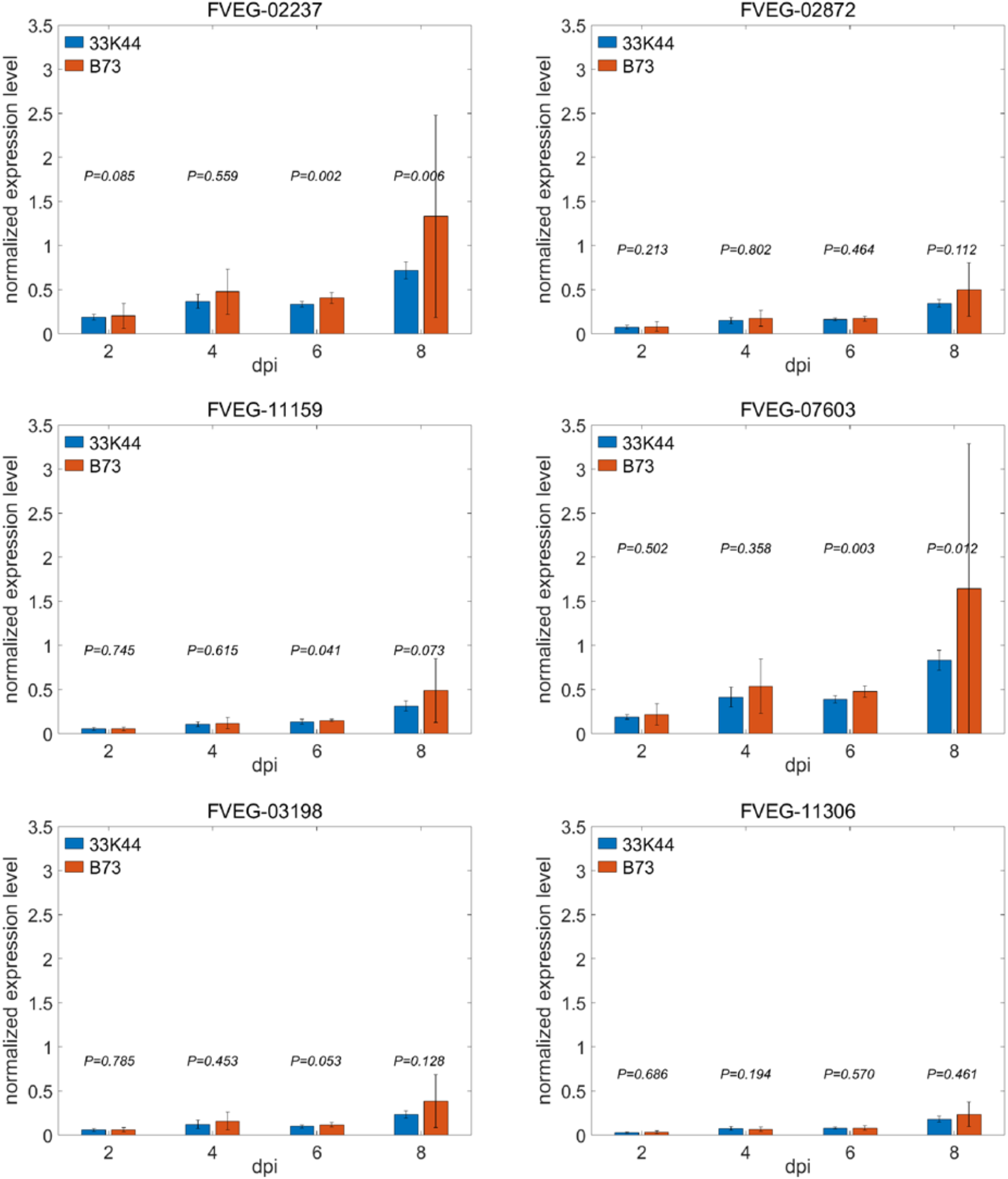
Transcription profile of 6 key Mediator genes in moderately resistant (B73) versus highly susceptible (33K44) maize seeds. *F. verticillioides* was inoculated on surface sterilized B73 and 33K44 seeds, and RNA-Seq was performed on 2, 4, 6, 8 days-post-inoculation (dpi) samples. We analyzed whether transcription of Mediator genes show difference in two maize seeds. *P value* for comparison was computed by the non-parametric Van der Waerden (VDW) test while not having any assumptions for normality or equality of variance between 33K44 and B73. A complete set of transcriptional comparisons for all *F. verticillioides* Mediator genes is shown in Fig. S1.

### Construction of Mediator complex deletion mutants

In an effort to test the biological functions of Mediator subunits in *F. verticillioides*, we generated knockout mutation of each gene via split-marker approach (Fig. S2). All transformants were analyzed by PCR and qPCR assays with corresponding primers to verify null mutation (Table S1). For 11 Mediator subunits, at least two knockout mutants with similar phenotypes were obtained (Table S2). As for the other 10 putative Mediator subunits (Table 1), we were not able to retrieve viable gene-deletion mutants after screening more than 200 transformants from at least four independent transformation experiments. This outcome led us to conclude that the null mutation of these subunits is deemed lethal in *F. verticillioides*. Significantly, these subunits are also recognized as essential genes for *S. cerevisiae* viability (33).

### Mediator complex play different roles in fungal development processes

When we tested vegetative growth in mutants, strains with Head module gene deletion (ΔFvMed18) and Kinase module gene deletion (ΔFvMed12, ΔFvMed13, ΔFvFcc1 and ΔFvFck1) exhibited significantly reduced growth rate compared to the wild-type progenitor on V8 agar (Fig. 2A and Table 2). We also compared colony morphology and growth rate of 11 Mediator mutants on 0.2×PDA and Myro agar plates (Table 2 and Fig. S3). Among these 11 mutants, 6 showed reduced conidiation by > 30% compared to the wild-type progenitor. Surprisingly, ΔFvMed9 and ΔFvMed5 exhibited significantly increased conidiation (Fig. 2B and Table 2). We also measured conidia germination rate in all Mediator subunit mutants and found that 9 mutants exhibited dramatic defect in conidia germination (Fig. 2C and Table 2). Interestingly, ΔFvMed18 conidia fail to germinate even after a 6.5-hour incubation on water agar (WA) plate. In addition, we observed that mycelial surface of some Mediator mutants exhibited different level of hydrophobicity compared with the wild-type progenitor. As showed in Fig. 2D, a 5-μL water droplet could persist for more than 1 h on WT and the majority of Mediator mutants. However, ΔFvMed18, ΔFvMed5, ΔFvMed16 and ΔFvMed12 exhibited different degree of reduced hydrophobicity (Fig. 2D). Taken together, these results suggested that Mediator complex is critical for fungal development processes in *F. verticillioides*, but each subunit likely have different principal functions.

**Fig. 2.**
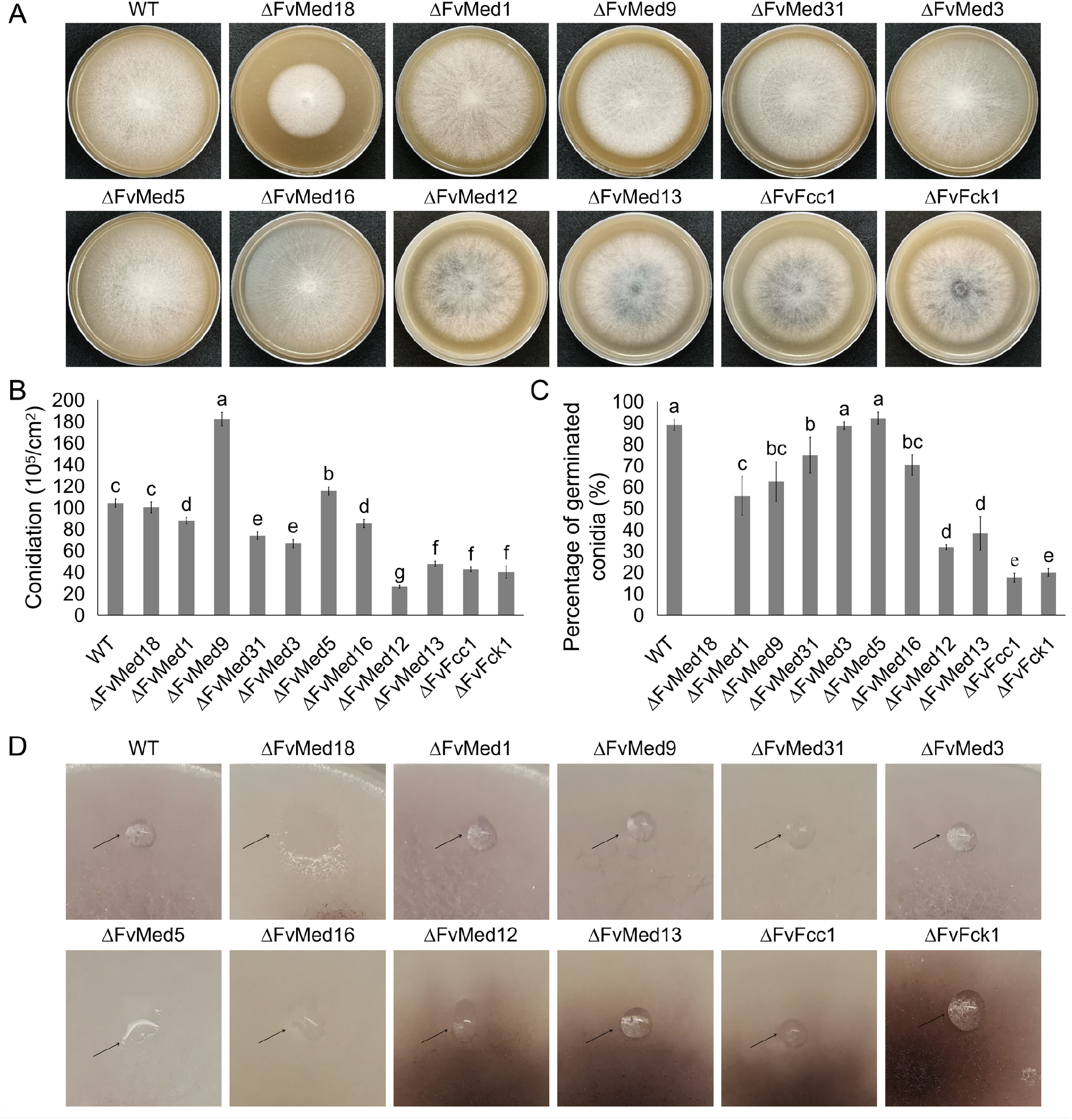
Phenotypes of the Mediator deletion mutants in vegetative growth, conidiation, conidial germination and hydrophobicity. (A) Colony morphology of wild-type and Mediator mutants grown on V8 plate at 28°C for 6 days. (B) Deletion of Mediator subunits on conidia production. Conidia of all strains were quantified with a hemacytometer after incubation on V8 plate for 6 days. (C) Conidia of each strain were cultivated in WA medium. After incubation at 28°C for 6.5 h, conidial germination of 200 conidia was examined. (D) Hydrophobicity assays of wild-type and all mutants. Bars denote standard errors from three repeated experiments. Values on the bars followed by the same letter are not significantly different at P = 0.05, according to Fisher’s LSD test.

**Table 2.** Hyphal growth, conidiation and conidial germination of the Mediator deletion mutants.

| Strain | Colony diameter (mm)<sup>a</sup> |  |  | Conidiation<br>(10<sup>5</sup>/cm<sup>2</sup>)<sup>b</sup> | Percentage of<br>germinated<br>conidia (%)<sup>c</sup> |
| --- | --- | --- | --- | --- | --- |
|  | V8 | PDA | Myro |  |  |
| WT | 80.0±1.4a | 63.0±0.8c | 66.8±0.9b | 104.1±3.9c | 89.0±2.6a |
| ΔFvMed18 | 44.7±1.2g | 38.2±0.8h | 29.5±1.4i | 100.2±5.0c | - |
| ΔFvMed1 | 78.0±1.4a | 59.8±1.1e | 54.5±0.8e | 87.8±2.9d | 55.8±9.1c |
| ΔFvMed9 | 66.3±0.9d | 59.0±1.2e | 61.3±1.4d | 182.2±6.2a | 62.5±9.3bc |
| ΔFvMed31 | 70.5±1.3c | 61.0±1.0de | 62.8±0.9cd | 73.7±3.7e | 74.8±8.4bc |
| ΔFvMed3 | 78.4±1.3a | 70.1±1.1a | 71.2±0.9a | 66.6±3.9e | 88.6±1.8a |
| ΔFvMed5 | 74.0±1.0b | 62.8±1.2cd | 68.0±1.4b | 115.4±3.8b | 92.2±2.8a |
| ΔFvMed16 | 75.0±1.0b | 65.2±0.9b | 64.3±1.3c | 85.2±3.8d | 70.2±4.7b |
| ΔFvMed12 | 64.1±1.2e | 54.8±1.2f | 40.5±1.2g | 26.4±1.4g | 31.8±1.2d |
| ΔFvMed13 | 69.0±1.1c | 60.0±0.9e | 43.6±1.4f | 47.6±2.4f | 38.2±7.9d |
| ΔFvFcc1 | 62.5±1.1ef | 53.4±1.2f | 35.1±0.8h | 42.5±2.1f | 17.6±2.1e |
| ΔFvFck1 | 61.5±0.9f | 50.7±1.4g | 41.7±0.8fg | 40.0±5.7f | 20.0±1.8e |

\*Means and standard deviations (SD) were calculated from three independent experiments. Values in table followed by the same letter are not significantly different ( $P < 0.05$ ).
<sup>a</sup>Colonies were cultured on V8, 0.2×PDA or Myro plates for 6 days at 28°C.
<sup>b</sup>Conidiation in V8 plate was examined after incubation for 6 days at 28°C.
<sup>c</sup>Conidial germination was assayed on Water Agar (WA) after 6.5 h of incubation at 28°C.

### Involvement of Mediator complex in plant virulence

To test the role Mediator subunits play in *F. verticillioides* virulence, we used maize seedling infection assay as previously described (34). After a 10-day incubation at room temperature, all mutants exhibited reduced virulence when compared to the wild-type progenitor. In particular, seedling rot symptoms caused by ΔFvMed18, ΔFvMed12, ΔFvMed13, ΔFvFcc1 and ΔFvFck1 were restricted around the inoculation sites (Fig. 3), suggesting that Head and Kinase modules in the Mediator complex play more crucial roles compared to Middle and Tail modules in fungal virulence.

**Fig. 3.**
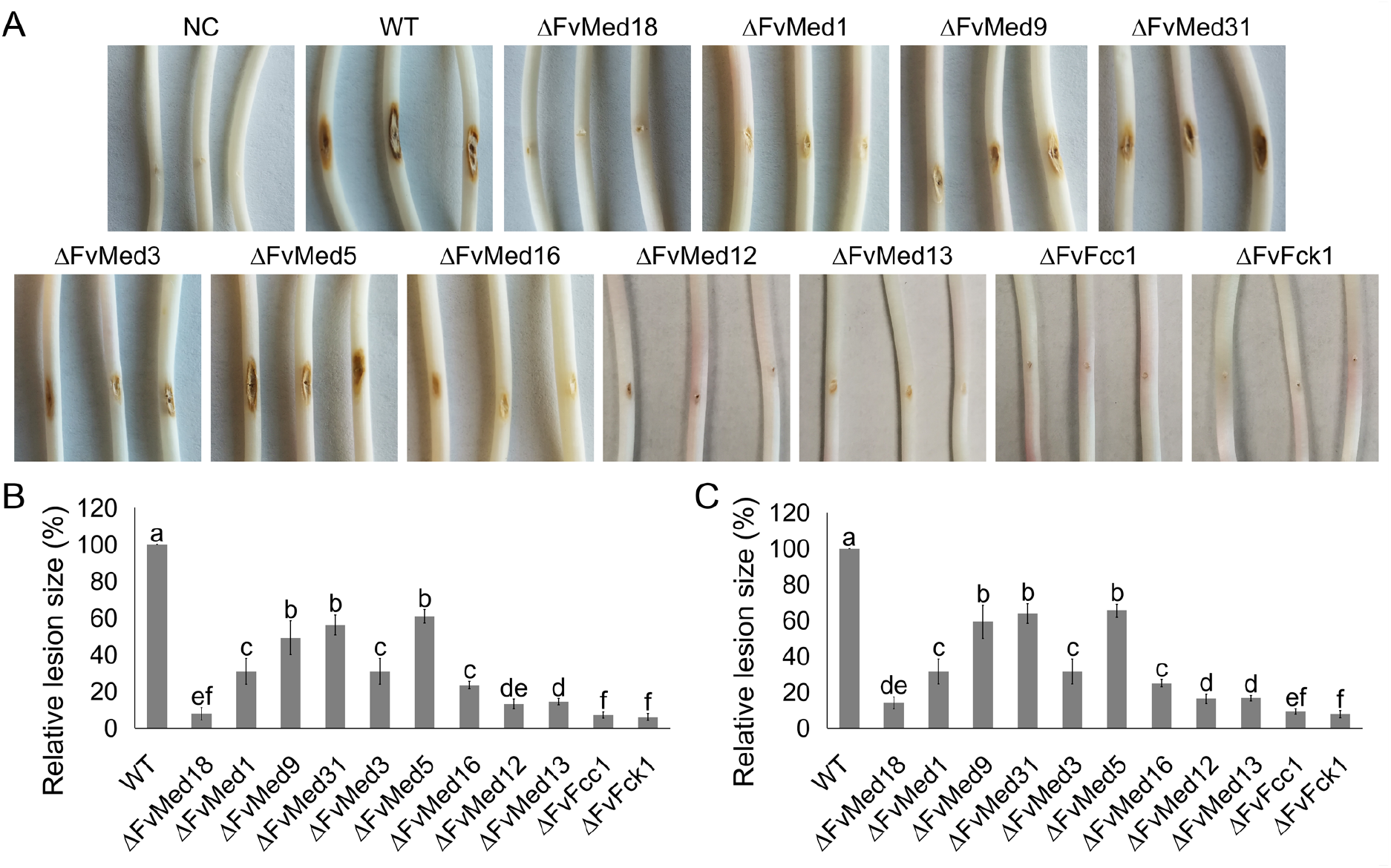
Impact of Mediator gene mutation in virulence when tested on maize seedlings. (A) Maize seedlings were grown in the dark and inoculated with conidia suspension from all strains. The images were taken after 10 days of incubation. Sterile water was used as negative control (NC). (B) The relative lesion size in maize seedling rot assays was measured by image J. (C) The relative lesion size was normalized with strain growth rate on V8 plate. Bars denote standard errors from three repeated experiments. Values on the bars followed by the same letter are not significantly different at P = 0.05, according to Fisher’s LSD test.

### Mediators show different levels of stress and fungicide sensitivity in *F. verticillioides*

We asked whether Mediator complex subunits play a role in regulating responses to environmental stress agents and subsequently tested all mutants against cell wall-damaging agent, oxidative and osmotic stresses, and fungicides. Our results showed that only 5 mutants (ΔFvMed18, ΔFvMed12, ΔFvMed13, ΔFvFcc1 and ΔFvFck1) exhibited slightly decreased sensitivity to cell wall-damaging agent Congo Red (Fig. 4 and Fig. S4). Overall, no sensitivity differences were observed among 11 Mediator mutants to osmotic stress in comparison with the wild-type progenitor in our experiments (data not shown).

**Fig. 4.**
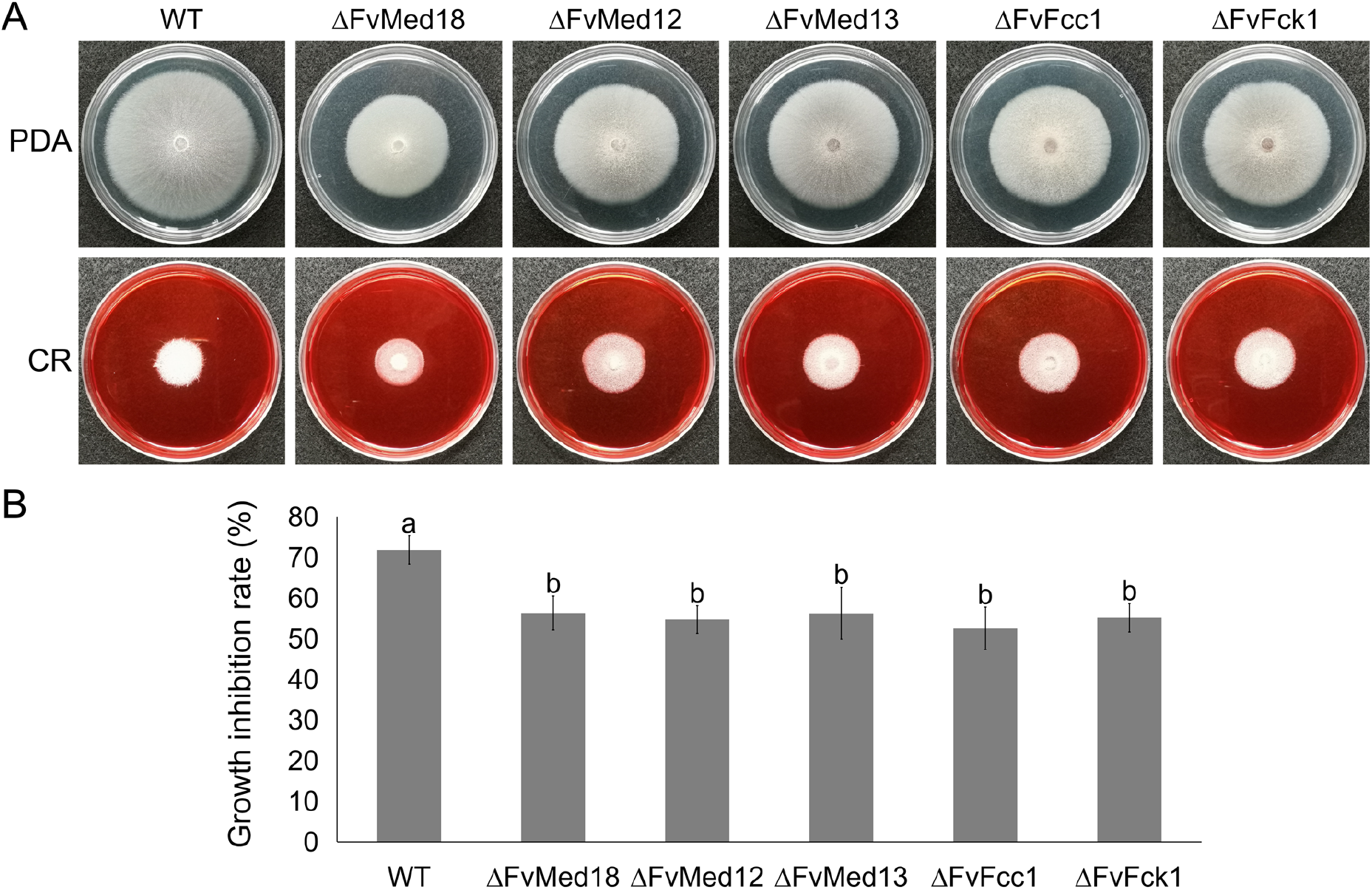
Gene disruption of Head and Kinase modules of Mediator increases the sensitivity towards cell-wall damage agent. (A) Growth phenotype of the wild type (WT) and mutants with Congo Red amended in 0.2xPDA plates after 6 days of incubation at 28°C. (B) Statistical analysis of the growth inhibition rate of each strain with Congo Red. Bars denote standard errors from three repeated experiments. Values on the bars followed by the same letter are not significantly different at P = 0.05, according to Fisher’s LSD test.

In human pathogens, Mediator has been shown to play an important role in azole fungicide resistance (35–39). Intriguingly, when we assayed the sensitivity of *F. verticillioides* Mediator mutants, deletion of Head and Kinase module subunits (ΔFvMed18, ΔFvMed12, ΔFvMed13, ΔFvFcc1 and ΔFvFck1) caused significant increase in ketoconazole and fluconazole resistance compared to the wild-type progenitor. However, no sensitivity difference was detected among other mutants (Table S3). These results demonstrated that Head and Kinase modules are negatively associated with azole resistance in *F. verticillioides*.

### Mediators are involved in carbon and fatty acid utilization

In earlier published studies, two homologs of Kinase module, CycC and Cdk8, were suggested as a link between ambient environment sensing and transcriptional regulation in *F. verticillioides* (26, 27). To further understand the roles of the Mediators in carbon source utilization, each mutant was cultured on minimal medium (MM) amended with glucose, fructose, lactose, starch or acetate as a sole carbon nutrient. As results showed, the vegetative growth of wild-type strain and all mutants were similar on MM amended with fructose, lactose and starch when compared to glucose (Fig. S5A). Interestingly, most of the mutants in ‘core Mediator’ exhibited reduced growth rate with acetate (except for ΔFvMed31 and ΔFvMed5). In contrast, mutants in Kinase module showed increased relative growth rate on MM amended with acetate as compared with glucose (Fig. 5). These results indicated that core Mediator plays contrasting role when compared to Kinase module in acetate utilization in *F. verticillioides*.

**Fig. 5.**
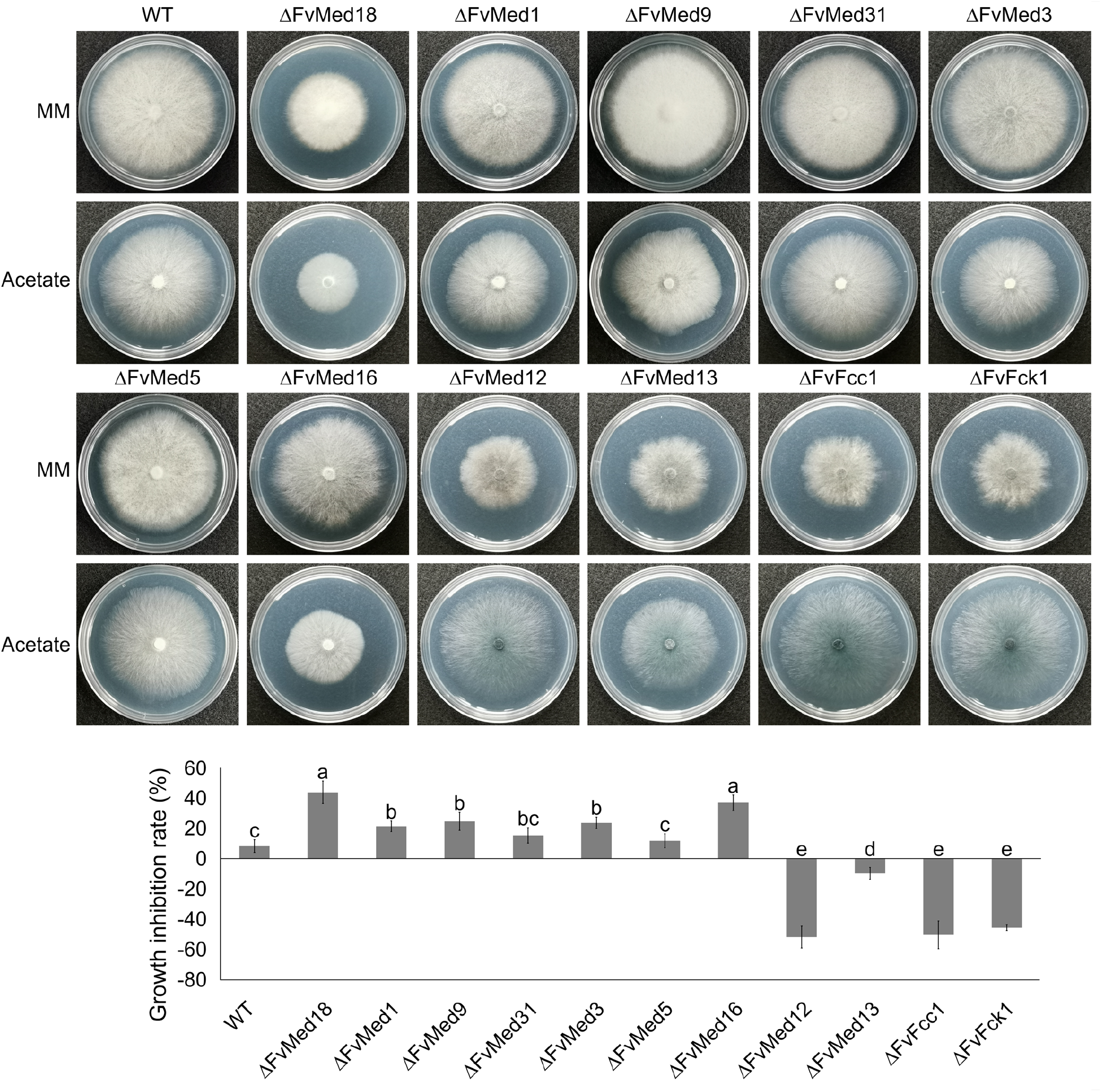
Mediator complex is involved in acetate utilization. (A) All strains were cultivated on MM with glucose or acetate, and incubated at 28°C for 6 days. (B) The growth inhibition rate of each strain was analyzed by comparing the growth with acetate against glucose. Bars denote standard errors from three repeated experiments. Values on the bars followed by the same letter are not significantly different at P = 0.05, according to Fisher’s LSD test.

We further tested carbon nutrient utilization using fatty acids. We observed that ΔFvMed18 exhibit reduced relative growth rate with butyrate sodium but increased relative growth rate with palmitic acid and corn oil compared with glucose (Fig. S5B). However, mutants in Kinase module showed increased relative growth rate with all fatty acids (excluding Med13 with butyrate sodium) tested in our study (Fig. S5B). These results suggested that subunits in Head and Kinase module are involved in fatty acids utilization, but their functions are discrepant.

### Mediator complexes play distinct roles in secondary metabolism

Fumonisin B_1_ (FB_1_) is the most harmful secondary metabolite produced by *F. verticillioides* (40). We assayed the FB_1_ production in Mediator mutants when they were incubated for 7 days on autoclaved corn kernels (26). After normalization with fungal ergosterol level in each sample set, we were able to confirm that FB_1_ production was significantly reduced in 10 Mediator mutants in comparison with the wild-type progenitor. One unexpected outcome was that ΔFvMed1 exhibited drastically higher FB_1_ biosynthesis than the wild type (Fig. 6A and Table S4). Consistently, the expression level of *PKS11 (FUM1)* was significantly up-regulated in ΔFvMed1 (Table S5). These results led us to hypothesize that FvMed1 functions as a negative regulator of FB_1_ biosynthesis in *F. verticillioides* while majority of other Mediator subunits serve as the positive regulator.

**Fig. 6.**
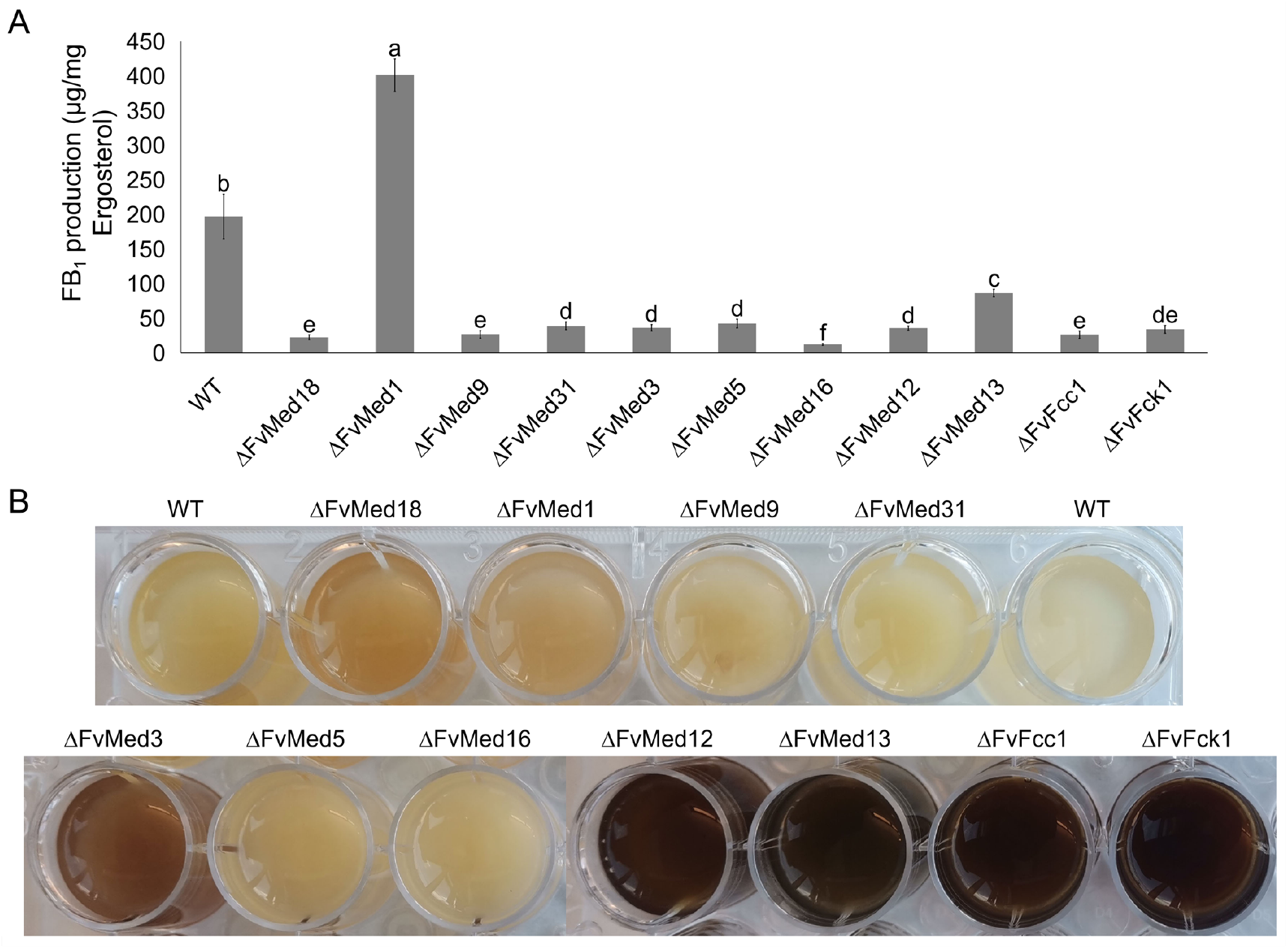
Roles of Mediator complex on FB_1_ and secondary metabolite biosynthesis. (A) FB_1_ production in the wild type (WT) and Mediator mutants grown on autoclaved kernels for 7 days at room temperature. FB_1_ production levels were normalized with ergosterol production in each strain. (B) Comparisons of secondary metabolite biosynthesis among the WT and Mediator mutants. Images were taken after all strains were grown on liquid Myro for 7 days. Bars denote standard errors from three repeated experiments. Values on the bars followed by the same letter are not significantly different at P = 0.05, according to Fisher’s LSD test.

To test the scope of Mediator complex subunit in regulating secondary metabolism in *F. verticillioides*, all mutants were cultured in YEPD (to promote vegetative growth) and Myro (to promote secondary metabolite production) liquid media for 3 days. As results showed, ΔFvMed3, ΔFvMed12, ΔFvMed13, ΔFvFcc1 and ΔFvFck1 exhibited higher levels of dark pigmentation in comparisons with the wild-type progenitor (Fig. 6B). Furthermore, qRT-PCR assays showed that the expression of gene associated with fusarubins biosynthesis, *PKS3*, is significant up-regulated in ΔFvMed18 and ΔFvFcc1, while deletion of ΔFvMed1 significant decrease *PKS3* expression (Table S5). Taken together, our results indicated that different Mediator modules and subunits regulate specific subset of genes involved in secondary metabolism in *F. verticillioides*.

### Subcellular localization analysis indicates functional variety of FvMed1

It is recognized that Mediator subunits are localized in cell nucleus and regulates gene transcription processes as an indispensable component of PIC (29, 41, 42). However, our FB_1_ assay results indicated that FvMed1 plays a unique role unlike other subunits under FB_1_ producing conditions. When we analyzed *in silico* putative Med1-interacting protein in *S. cerevisiae*, majority of the published and predicted interacting proteins were nuclear proteins, *e.g.*, subunits of Mediator complexes and histone proteins. However, we learned that certain non-nuclear proteins, e.g. cytoplasmic proteins, mitochondrial proteins and endoplasmic reticulum proteins, were identified as Med1-interacting proteins in *S. cerevisiae*. This raised a question whether Med1 can translocate outside the nucleus in response to certain ambient signals or conditions. In order to test this idea, we generated a FvMed1-GFP fusion construct with its native promoter and transformed it into the wild-type protoplast. All transformants were verified by PCR assay and observed under fluorescence microscope. Under FB_1_ non-producing condition, we observed weak GFP signals likely in the nucleus but clearly not in vacuoles in each septate mycelial cell (Fig. 7A). Under FB_1_-producing condition, although Med1-GFP could be found in the nucleus, we also observed multiple punctate and rounded structures in the cytoplasm (Fig. 7B - D). Significantly, we did not observe GFP signals localizing to vacuoles as well as a large number of lipid bodies, which are commonly observed when F. verticillioides is grown in secondary metabolite-inducing media. Also, that fact that the GFP signal was not diffused throughout the cytoplasm suggests that FvMed1 translocates to a defined cellular organelle when it moves out of the nucleus. When assayed for *FvMed1* expression in YEPD and Myro cultures, the transcript level of *FvMed1* showed no difference between two media (data not shown), suggesting that *FvMed1* may be constitutively expressed in different growth conditions. These results showed that FvMed1 proteins were transported to the cytoplasm in response to FB_1_ promoting signals.

**Fig. 7.**
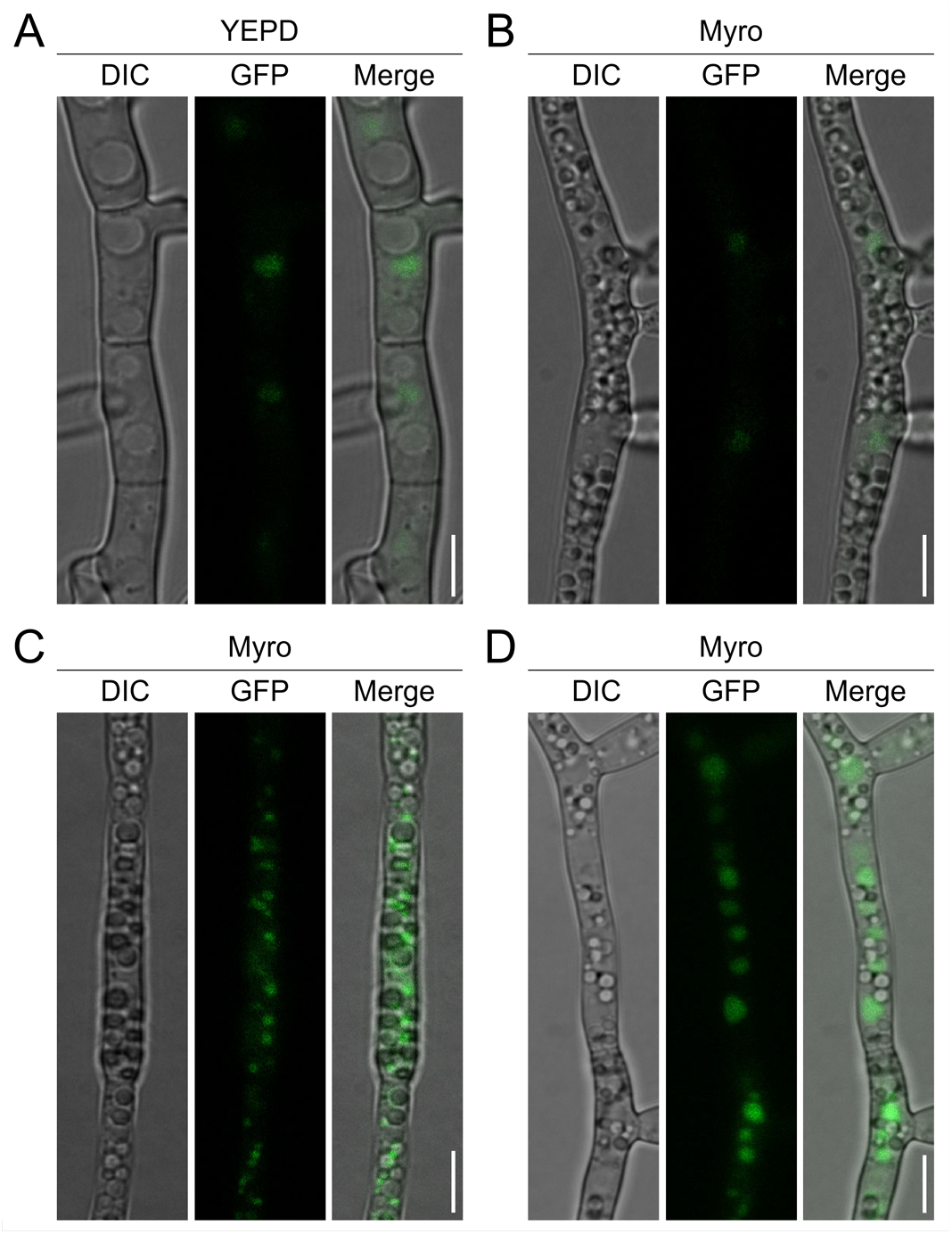
Subcellular localization of Med1-GFP in *F. verticillioides* mycelial cells grown in (A) FB_1_-non-inducing medium YEP and (B~D) FB_1_-inducing medium Myro. We inoculated *F. verticillioides* Med1-GFP conidia in two cultures and took images after 2-day incubation. GFP expression was observedunder an Olympus BX51 microscope and the images were processed by ImageJ. Bar = 10 μm.

## DISCUSSION

Mediator is a conserved protein complex that performs indispensable cellular functions as a regulator of gene transcription. Here, we identified 21 genes encoding putative *F. verticillioides* Mediator homologs by using 25 *S. cerevisiae* Mediator subunits as queries. Although the modular organization of Mediator complexes are predicted to be similar between *S. cerevisiae* and *F. verticillioides*, the biological functions of certain subunits show significant differences in *F. verticillioides*. For instance, the gene encoding *Med15* resides as a single copy locus in *S. cerevisiae*, but the homolog in *F. verticillioides* seems to have expanded to 11 copies of telomere-associated genes (*TLO*s). We are making this prediction base on previous report that showed *S. cerevisiae Med2* being replaced by a subset of *TLOs* in *C. albicans* (42). Furthermore, when we studied 11 viable gene knockout mutants, we observed some distinct phenotypic outcomes in different mutants.

It is perceived that Mediator is important for fungal development processes, secondary metabolism and plant infection. Intriguingly, we observed phenotypic variations when comparing between modules and subunits, indicating that certain modules or subunits of Mediator complex preferentially control the expression of different subset of target genes, thus regulating specific developmental processes. Disruption of a Kinase module exhibited defects in growth and conidia production in *F. graminearum* (28, 29). Another study showed that *CaMed7* is required for hyphal development (16). In our study, 5 Mediator mutants displayed significant defect in vegetative growth, including mutants in Head and Kinase modules. In addition, while the conidia production in ΔFvMed18 was similar to the wild-type strain, the conidial germination was conspicuously suppressed. Furthermore, we observed that conidia produced in ΔFvMed9 and ΔFvMed5 were significantly increased compared to the wild-type progenitor. These results indicated that each module or subunit of Mediator have unique functions during fungal development processes in *F. verticillioides*.

In *Cryptococcus neoformans, Cdk8* and *Ssn8* deletion mutants showed increased sensitivity to cell wall and oxidative stress (44). The authors provided further evidence that this outcome was due to mitochondria morphology defect and the resulting transcriptional regulation defects were associated with oxidation-reduction mechanism. In our study, we observed that mutants in Head and Kinase modules exhibited reduced tolerance to cell wall damage agent. However, no significant difference was found in wild-type strain and all Mediator mutants in regard to oxidative stress agents (data not shown). These results suggested that Mediator is involved in regulating environment stress responses, but there may be species-specific functions against various stress agents. Elucidating the mechanism in which these processes are disrupted in *F. verticillioides* may require further study.

Glucose is the most suitable carbon source for eukaryotes, especially fungi. However, fungi can utilize a wide array of carbon sources, even the most complex carbohydrates, to survive and thrive in the most challenging environmental conditions. Previous research showed that certain subunits of Mediator complex are involved in glucose metabolism and carbon source utilization (14, 44–46). Our study showed how each Mediator mutant can differ in its ability to utilize different carbon sources. While most of the mutants in the core Mediator showed defects in using acetate as sole carbon source, mutants in Kinase module exhibited increased growth rate in MM amended with acetate when compared to glucose. Surprisingly, these results do not agree with those observed in *C. neoformans* ΔCdk8 and ΔSsn8 mutants (44), indicating that Mediator has species-specific functions in regulating genes involved in glucose metabolism.

A number of studies have demonstrated the importance role of Mediator in virulence. In *Candida glabrata*, the Med2 subunit was shown to control the Epa adhesins biosynthesis, with the mutant displaying elevated adherence to epithelial cells (37). In *C. albicans*, Med13 and Med31 subunits can activate genes associated with adhesion thereby stimulating fungal infection (47). Here, we observed that all Mediator mutants in *F. verticillioides* exhibited attenuated disease symptoms on maize seedling, despite each Mediator mutant showing varying reduced virulence levels. Additionally, our results are not consistent with those observed in *C. albicans*, further supporting the divergence of function within Mediator despite their conserved roles across eukaryotes.

Our effort to effectively control fungal diseases in crop production faces persistent difficulty due to the limitation in antifungal agent availability. More seriously, this difficulty is exacerbated due to the emergence of resistant pathogen population in the field. Many studies showed that Mediator plays an important role in azoles resistance by regulating transcriptional level of efflux pump genes. In *C. glabrata*, the expression of efflux pump genes *CgCDR1* and *CgCDR2* were significantly decreased in *CgMed15A* mutant, which resulted in increased sensibility to azole fungicides (36, 48). In addition, deletion of *CgMed2* also resulted in the reduction of *CgCDR1* expression and sensitivity to azoles (37). However, *CgMed1* deletion mutant did not exhibit any changes to azole sensitivity and *CDR2* expression (48). In our study, we tested the sensitivity of wild-type and all Mediator mutants against two azole fungicides ketoconazole and fluconazole. We observed that mutants in Head and Kinase module exhibited increased drug resistance compared to the wild-type and other mutant strains. However, qPCR assays showed that the expression of efflux pump genes in these mutants showed no statistical difference in comparison to the wild-type progenitor and other mutants. Intriguingly, recent studies showed that the deletion of *Ssn3* or *Ssn8* could increase azole resistance, which is associated with chromatin remodeling (49, 50). Thus, we hypothesize that the increased drug resistance in *F. verticillioides* Mediator mutants share similar mechanism to that in *C. albicans*.

In our study, FB_1_ biosynthesis was significantly reduced in majority of the Mediator mutant except ΔFvMed1 and ΔFvMed9. Similar results were observed in *F. graminearum* (28, 29). Surprisingly, we found that FB_1_ biosynthesis in ΔFvMed1 was about 2-fold higher than wild-type progenitor. We questioned whether FvMed1 plays different roles in other secondary metabolism compared to other Mediator subunits, and analyzed the expression of all 15 *PKS* genes in ΔFvMed1 under toxin-inducing conditions. We learned that the expression of *PKS11 (FUM1)* was significantly higher in ΔFvMed1. Additionally, we observed that the expression of other *PKS* genes showed significantly difference among 4 Mediator mutants tested (Table S5). These results implied that Mediator complex is critical for secondary metabolism in phytopathogen, but each subunit played unique roles and regulated specific subsets of genes.

Although many reports focused on the functions of Mediator subunits in the nucleus and as a component to RNA Pol II, an earlier study revealed their possible roles in the cytoplasm (51). In lung cancer cells, Med12 was found in both the nucleus and the cytoplasm. When localized in the cytoplasm, Med12 participated in TGF-beta signaling regulation via binding with TGF-beta receptor 2, thereby affecting its glycosylation and cell membrane expression (51). We hypothesize that FvMed1 can translocate to the cytoplasm under FB_1_-producing conditions and perhaps perform its unique function as a cytoplasmic protein. Our result raised a possibility that FvMed1 can control yet-to-be determined cytoplasmic signaling pathways associated with FB_1_ biosynthesis. Although the potential cytoplasmic FvMed1-interacting proteins remains unknown, this outcome provides us with motivation to expand our efforts to gain novel insight into the cytoplasmic function of Mediator subunits in filamentous fungi.

## MATERIALS AND METHODS

### Strains and culture assays

*F. verticillioides* wild-type strain M3125 was used for the construction of the derived mutants used in this study (34). All strains used in this study were grown and evaluated at 28°C on V8, 0.2×PDA and Myro agar plates for mycelial growth (34). For conidia gemination assay, conidia (10^6^) grown on V8 plates for 6 days were harvested and subsequently cultivated in water agar plates for 6.5 h at 28°C in dark. For stress response assays, 5-mm diameter mycelial plugs taken from a 6-day-old colony of wild-type and all mutant strains grown on O.2×PDA were inoculated on O.2×PDA plate containing 0.05% Congo red (w/v) or NaCl (1.2 M). Carbon source utilization assays were performed on minimal medium (MM) supplemented with different carbon sources at concentrations of 1% (w/v). All experiments were repeated three times independently.

### RNA-Seq data analysis

RNA-seq data were obtained through sequencing platform, Illumina HiSeq 2500, from Texas A&M AgriLife Genomics and Bioinformatics Services (College Station, Texas). The platform processed fourteen independent samples composed of seven biological replicates and seven technical replicates at four distinct time points (2 dpi, 4 dpi, 6 dpi, and 8 dpi). The sequencing data were generated as paired-end reads with 125-bp for two different maize kernels (hybrid 33k44 vs. inbred B73), where *F. verticillioides* inoculated. We first aligned and quantified the RNA-seq by TopHat2 (51) along with HTSeq (53). Subsequently, we filtered scarcely expressed genes out and normalized the remaining quantity by two steps, TPM (54) and TMM (55). Our dataset is publicly available at NCBI Gene Expression Omnibus (Accession GSE146208).

For those selected genes, we analyzed whether their expression levels were directly/indirectly associated with the virulence by comparing expression levels of 33K44 and those of B73 at all the time points. *P value* for comparison was computed by the non-parametric Van der Waerden (VDW) test while not having any assumptions for normality or equality of variance between 33K44 and B73. The bar plots of their mean expression levels with *P values* for the comparison are shown in Fig. 1 and Fig. S1.

### Strains construction

All gene deletion mutants were constructed using our standard split-marker gene knock out protocol as previously described (34). Hygromycin B phosphotransferase gene *(HPH)* was split into two fragments *PH* (929 bp) and *HP* (765 bp) and fused with left and right flanking sequence fragments of each targeted gene, respectively (Fig. S2). As the result of this homologous recombination, the open reading frame (ORF) of each gene was replaced with *HPH* gene. We confirmed targeted gene replacement by PCR, and further tested the absence of gene transcription by qPCR. FvMed1-GFP fusion cassette was constructed and transformed into *F. verticillioides* following our standard protocol described previously (34). All transformants were verified by PCR and observed under an Olympus BX51 microscope (Olympus America, Melville, NY, USA). A detailed description of features used for imaging fluorescence with this microscope and the image processing has been described previously (34).

### Sensitivity assay against ketoconazole and fluconazole

To determine the sensitivity to ketoconazole and fluconazole, a mycelial plug of all strains from a 6-day-old 0.2×PDA plate was transferred to the center of 0.2×PDA plates amended with ketoconazole and fluconazole at different concentrations. Three replicates for all concentrations were tested. After incubation at 28°C for 6 days, the colony diameter in all plates were measured. For each plate, the average colony diameter was used to calculate the fungicide concentration that resulted in 50% mycelial growth inhibition (EC50). The EC50 values were calculated with the Data Processing System (DPS) software. Ketoconazole was added to 0.2×PDA plates at 0, 0.01, 0.04, 0.2, 1.0 and 5.0 μg/mL. Fluconazole was added to 0.2×PDA plates at 0, 0.2, 1.0, 5.0, 10.0 and 50.0 μg/mL. The assays were repeated three time.

### RNA preparation and qPCR assays for Mediator subunit genes

To analyze relative gene expression of Mediator subunits, fungal mycelia were harvested from 3-days-old YEPD or 3-days-old Myro cultures and used for RNA isolation with the GeneJet RNA Purification Kit (Thermo Scientific). First-strand cDNA was synthesis by Verso cDNA synthesis kit (Thermo Scientific) and subsequently used for qPCR assays with DyNAmo ColorFlash SYBR Green qPCR Kit (Thermo Scientific). The β-tubulin gene of *F. verticillioides* was used as the internal control. The relative quantification of all genes was calculated with the 2^-ΔΔCt^ method (56). All qPCR assays were conducted with a StepOne plus realtime PCR system. All results were calculated with the data from three biological replicates.

### Fumonisin B_1_ assay and Maize seedling infection assay

To determine FB_1_ and ergosterol production, we followed the standard method described previously (34, 57). FB_1_ levels were normalized with ergosterol levels. All results were calculated with the data from three biological replicates. Stalk rot virulence assays was conducted using silver queen hybrid seeds as previously described (34). After growth on V8 plates for 6 days, conidia of each mutant were collected and re-suspended in sterile distilled water to a concentration of 1×10^7^ ml.^-1^. Seedlings were inoculated with 10 μL conidia suspension. After 10 days inoculation in the dark room, the infected seedlings were recorded and imaged. For all mutants, 15 seedlings were inoculated.

## ACKNOWLEDGMENTS

The research was supported by Texas A&M T3: Triads for Transformation Program and the China Scholarship Council State Scholarship Fund (201906850058). The authors declare no conflicts of interest.

---

**Table S1.**
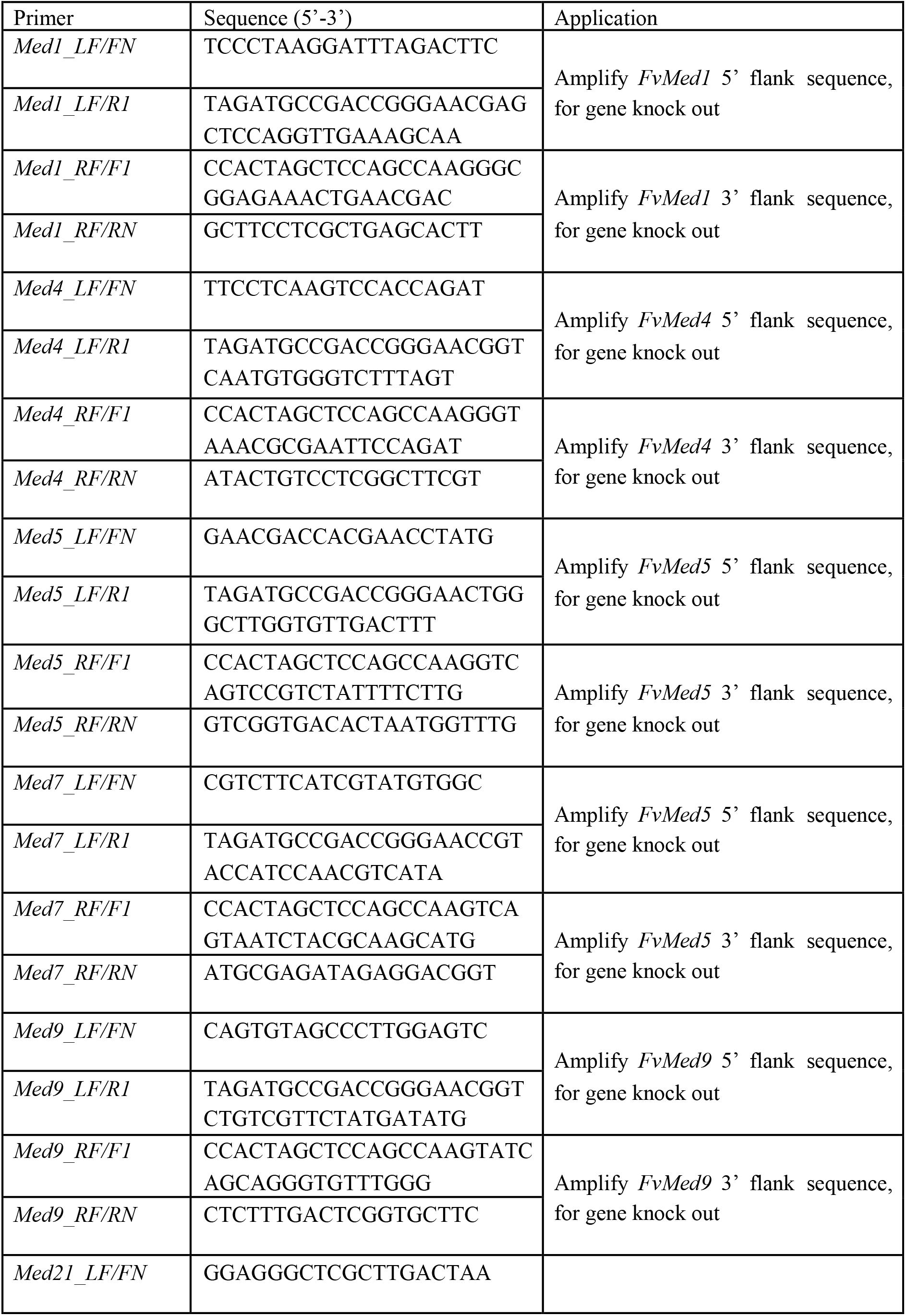
Primers used in this study.

**Table S2.** Wild-type and mutant strains of *F. verticillioides* used in this study.

| Strain | Genotype description | Reference |
| --- | --- | --- |
| M3125 | Wild-type strain of *F. verticillioides* | Lab preservation |
| ΔFvMed18 | *FvMed18* deletion mutant of M3125 | This study |
| ΔFvMed1 | *FvMed1* deletion mutant of M3125 | This study |
| ΔFvMed9 | *FvMed9* deletion mutant of M3125 | This study |
| ΔFvMed31 | *FvMed31* deletion mutant of M3125 | This study |
| ΔFvMed3 | *FvMed3* deletion mutant of M3125 | This study |
| ΔFvMed5 | *FvMed5* deletion mutant of M3125 | This study |
| ΔFvMed16 | *FvMed16* deletion mutant of M3125 | This study |
| ΔFvMed12 | *FvMed12* deletion mutant of M3125 | This study |
| ΔFvMed13 | *FvMed13* deletion mutant of M3125 | This study |
| ΔFvFcc1 | *FvFcc1* deletion mutant of M3125 | Shim and<br>Woloshuk, 2001<br>and this study |
| ΔFvFck1 | *FvFck1* deletion mutant of M3125 | Bluhm and<br>Woloshuk, 2005<br>and this study |

**Table S3.** Sensitivity of *F. verticillioides* to ketoconazole and fluconazole.

| Strain | EC<sub>50</sub> (µg/mL) |  |
| --- | --- | --- |
|  | Ketoconazole | Fluconazole |
| WT | 1.41±0.11c | 1.32±0.10c |
| ΔFvMed18 | 2.76±0.30ab | 4.09±0.86a |
| ΔFvMed1 | 1.48±0.11c | 1.38±0.05c |
| ΔFvMed9 | 1.43±0.11c | 1.34±0.03c |
| ΔFvMed31 | 1.32±0.16c | 1.29±0.16c |
| ΔFvMed3 | 1.45±0.15c | 1.36±0.10c |
| ΔFvMed5 | 1.55±0.09c | 1.45±0.05c |
| ΔFvMed16 | 1.36±0.32c | 1.28±0.16c |
| ΔFvMed12 | 3.08±0.24a | 2.17±0.21b |
| ΔFvMed13 | 2.65±0.08b | 2.06±0.06b |
| ΔFvFcc1 | 2.43±0.19b | 2.10±0.07b |
| ΔFvFck1 | 2.84±0.22ab | 2.17±0.18b |

\*Means and standard deviations (SD) were calculated from three independent experiments.
Values in table followed by the same letter are not significantly different ( $P < 0.05$ ).
EC<sub>50</sub>: the fungicide concentration resulting in 50% inhibition of mycelial growth.

**Table S4.** Fumonisin B_1_ (FB_1_) and ergosterolproduction of the wild-type and Mediator deletion mutants.

|  | FB<sub>1</sub> (μg/L)<sup>a</sup> | Ergosterol (mg/L)<sup>b</sup> | FB<sub>1</sub>/Ergosterol (μg/mg)<sup>c</sup> |
| --- | --- | --- | --- |
| WT | 73915.6±12127.6a | 375.0±49.8a | 197.1±32.3b |
| ΔFvMed18 | 4797.6±712.0e | 213.1±29.4b | 22.5±3.4d |
| ΔFvMed1 | 84499.1±4949.8a | 210.6±9.9b | 401.3±23.5a |
| ΔFvMed9 | 6719.5±799.1d | 248.3±27.5b | 27.1±7.1d |
| ΔFvMed31 | 9551.1±1822.6c | 234.3±41.0b | 39.3±7.5d |
| ΔFvMed3 | 13488.1±1790.0b | 370.2±53.6a | 36.4±4.8d |
| ΔFvMed5 | 10845.0±1175.8bc | 254.9±24.7b | 42.5±6.5d |
| ΔFvMed16 | 3270.1±353.0f | 264.5±47.0b | 12.4±1.3d |
| ΔFvMed12 | 2475.8±196.8f | 68.7±14.3cd | 36.0±2.9d |
| ΔFvMed13 | 6349.3±407.3d | 73.5±16.7cd | 86.4±5.5c |
| ΔFvFcc1 | 1636.1±327.9g | 62.0±8.8d | 26.4±5.3d |
| ΔFvFck1 | 2859.8±583.3f | 83.5±6.0c | 34.2±7.0d |

\*Means and standard deviations (SD) were calculated from three independent experiments. Values in table followed by the same letter are not significantly different ( $P < 0.05$ ).
<sup>a</sup>FB<sub>1</sub> production in wild-type and mutants grown on autoclaved kernels for 7 days at room temperature.
<sup>b</sup>Ergosterol production in wild-type and mutants grown on autoclaved kernels for 7 days at room temperature.
<sup>c</sup>FB<sub>1</sub> levels were normalized with fungal ergosterol levels.

**Table S5.** Relative expression level of 15 *PKS* genes in wild-type and Mediator mutants.

| Gene | Relative expression level |  |  |  |  |
| --- | --- | --- | --- | --- | --- |
| | WT | $\Delta FvMed1$ | $\Delta FvMed3$ | $\Delta FvMed18$ | $\Delta FvFcc1$ |
| *PKS1* | 1 | 2.55±0.33 | ND | ND | 8.78±2.52 |
| *PKS2* | 1 | 0.97±0.10 | 0.98±0.22 | 0.64±0.06 | 3.74±0.37 |
| *PKS3* | 1 | 0.36±0.03 | 1.93±0.77 | 8.66±0.84 | 31.17±3.65 |
| *PKS4* | 1 | 8.41±1.12 | 0.24±0.11 | 0.29±0.11 | 9.92±1.52 |
| *PKS5* | 1 | 5.18±0.78 | 1.00±0.15 | ND | 1.14±0.17 |
| *PKS6* | 1 | 2.30±0.21 | 1.85±0.31 | 0.02±0.01 | 3.70±0.26 |
| *PKS7* | 1 | 0.47±0.08 | 2.44±0.36 | 3.18±0.45 | 0.63±0.16 |
| *PKS8* | 1 | 3.70±0.35 | 11.03±1.96 | 0.03±0.003 | 0.05±0.01 |
| *PKS9* | 1 | 0.31±0.05 | 0.62±0.04 | 0.56±0.06 | 0.27±0.02 |
| *PKS10* | 1 | 0.14±0.04 | 2.34±0.34 | 0.07±0.04 | 0.58±0.04 |
| *PKS11* | 1 | 2.49±0.27 | 0.62±0.2 | 0.04±0.01 | 0.06±0.01 |
| *PKS12* | ND | ND | ND | ND | ND |
| *PKS13* | 1 | 1.13±0.07 | 2.25±0.13 | 1.21±0.07 | 1.80±0.11 |
| *PKS14* | 1 | 0.06±0.01 | ND | 0.24±0.06 | 0.08±0.01 |
| *PKS15* | 1 | 0.66±0.2 | 0.23±0.07 | 65.78±21.22 | 1.42±0.33 |

Mycelia of each mutants were incubated in myro liquid medium for 3 days at 28°C. Each gene expression were normalized with β-tubulin expression level. Gene expressions were calculated using $2^{-\Delta\Delta Ct}$ . Gene expression level of wild-type strain was standardized to 1.0. Three replicates were performed for each test. ND: not detected.

**Fig. S1.**
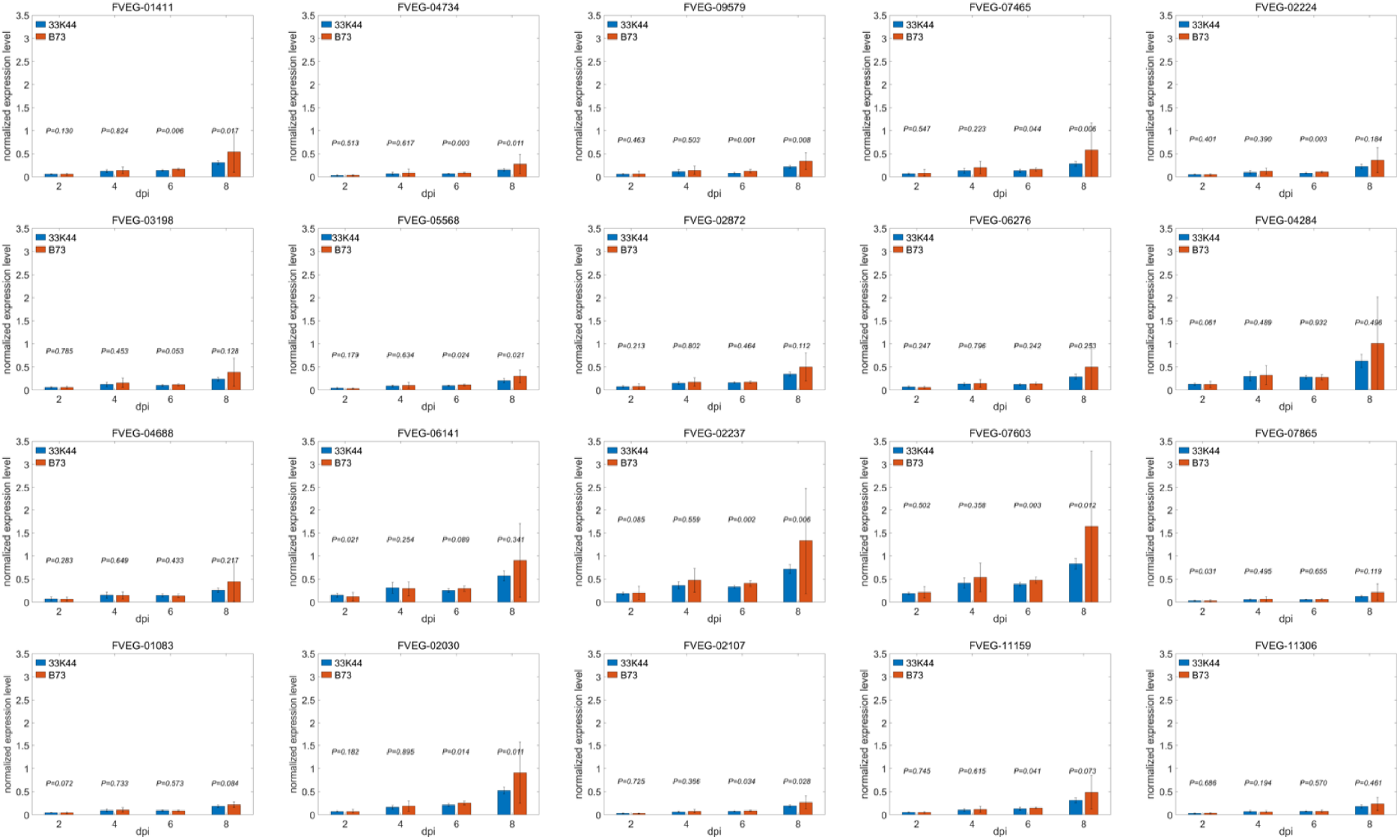
Assessment of the RNA-Seq Data Set of *F. verticillioides* Mediator genes under different conditions. The Mediator genes expression were normalized with β-tubulin expression. Error bars indicate SD of 7 biological replicates.

**Fig S2.**
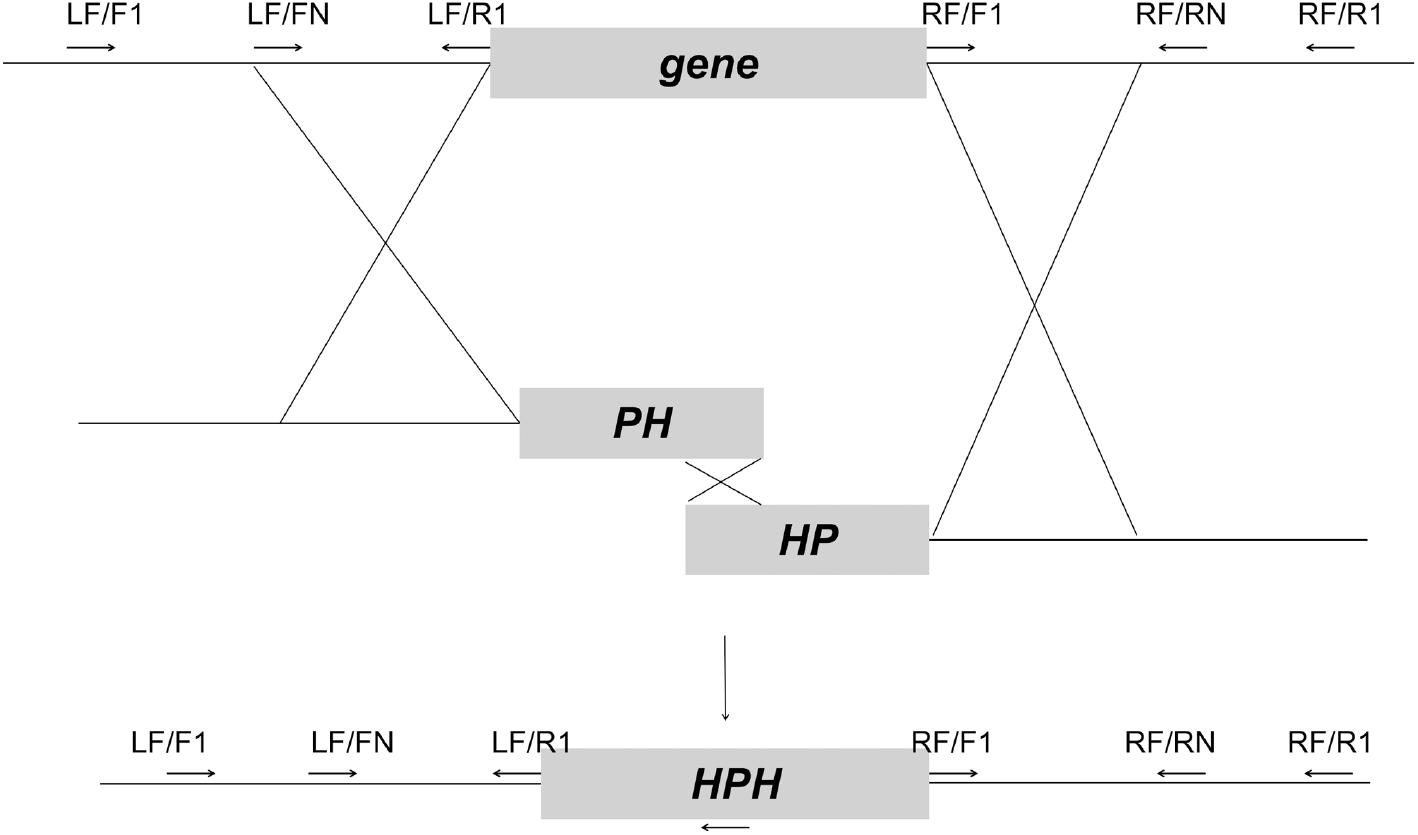
Schematic description of split marker approach for constructing Mediator deletion mutants in *F. verticillioides*.

**Fig S3.**
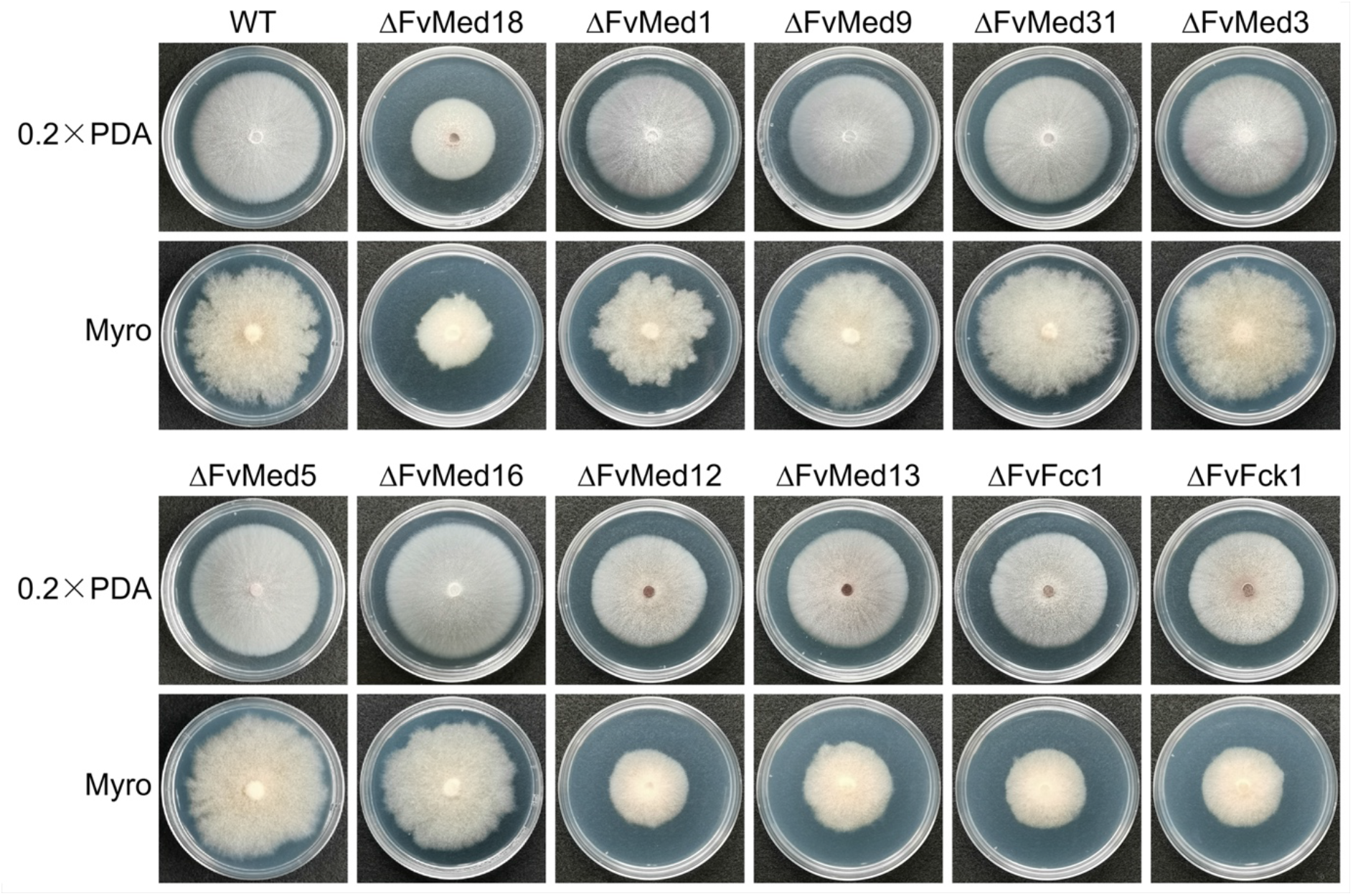
Colony morphology of wild-type and Mediator mutants grown on 0.2×PDA and Myro agar plate at 25°C for 6 days

**Fig S4.**
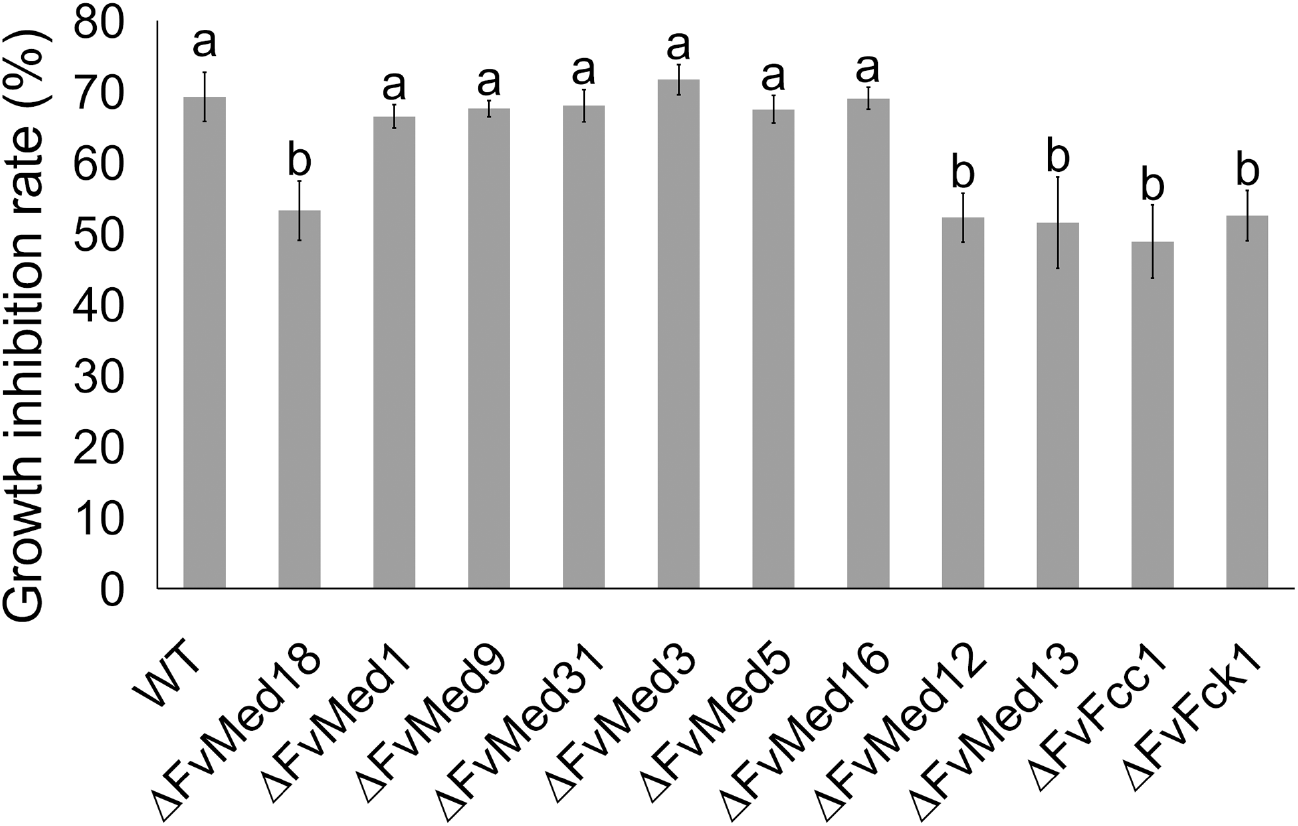
Deletion of Mediator subunits on cell-wall damage agent response in *F. verticillioides*

**Fig S5.**
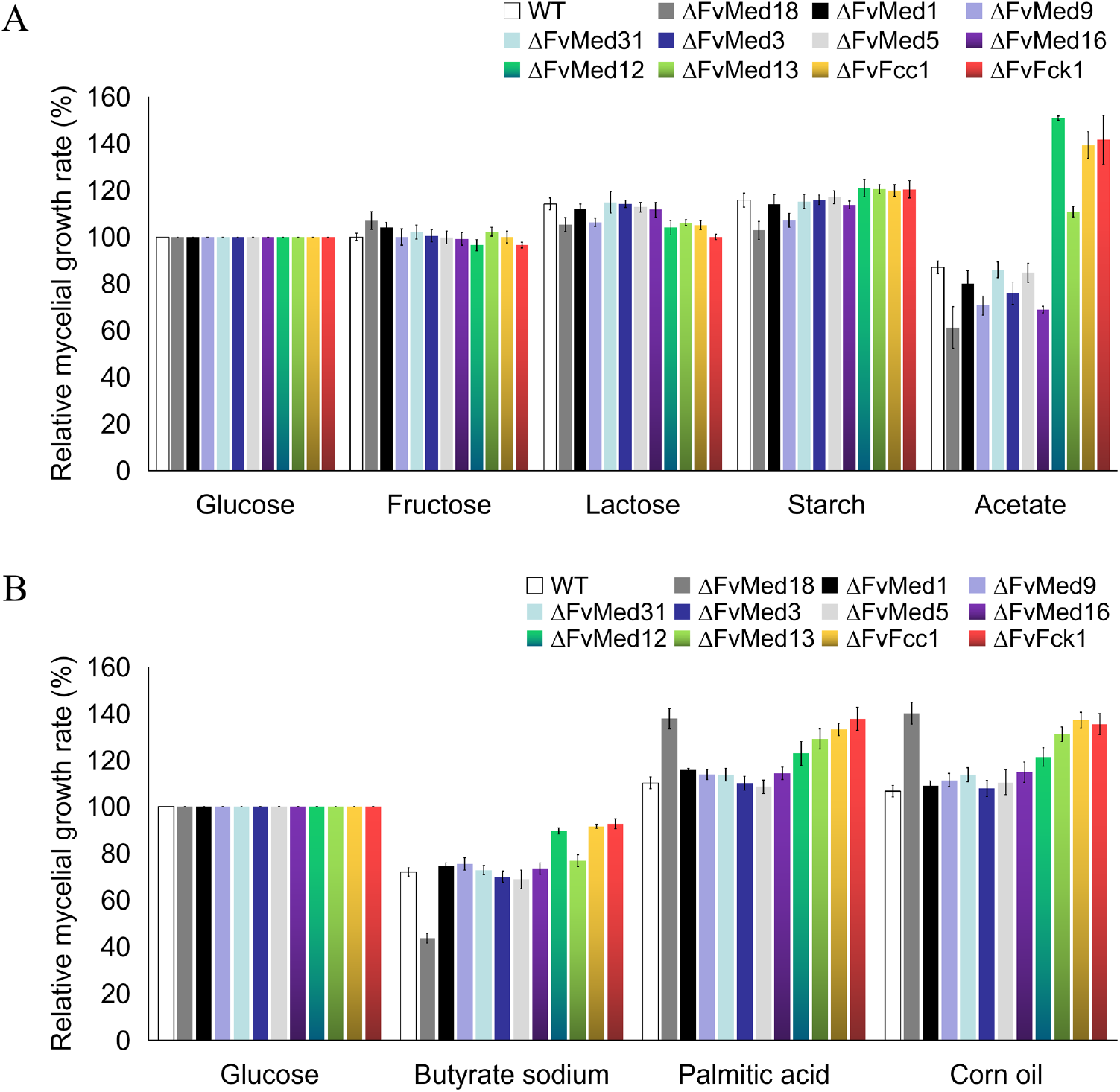
Relative growth rates of wild-type and Mediator mutants on various carbon sources. Values on the bars followed by the same letter are not significantly different at P = 0.05, according to Fisher’s LSD test

